# Combining experiments and in silico modeling to infer the role of adhesion and proliferation on the collective dynamics of cells

**DOI:** 10.1101/2021.03.29.437400

**Authors:** Hygor P. M. Melo, F. Raquel Maia, André S. Nunes, Rui L. Reis, Joaquim M. Oliveira, Nuno A. M. Araújo

**Affiliations:** Centro de Física Teórica e Computacional, Faculdade de Ciências, Universidade de Lisboa, 1749–016 Lisboa, Portugal; 3B’s Research Group, I3Bs - Research Institute on Biomaterials, Biodegradables and Biomimetics of University of Minho, Headquarters of the European Institute of Excellence on Tissue Engineering and Regenerative Medicine, AvePark, Parque de Ciência e Tecnologia, Zona Industrial da Gandra, 4805-017 Barco, Guimarães, Portugal; ICVS/3B’s – PT Government Associate Laboratory, Braga/Guimarães, Portugal; Departamento de Física, Faculdade de Ciências, Universidade de Lisboa, 1749–016 Lisboa, Portugal

## Abstract

The collective dynamics of cells on surfaces and interfaces poses technological and theoretical challenges in the study of morphogenesis, tissue engineering, and cancer. Different mechanisms are at play, including, cell-cell adhesion, cell motility, and proliferation. However, the relative importance of each one is elusive. Here, experiments with a culture of glioblastoma multiforme cells on a substrate are combined with in silico modeling to infer the rate of each mechanism. By parametrizing these rates, the time-dependence of the spatial correlation observed experimentally is reproduced. The obtained results suggest a reduction in cell-cell adhesion with the density of cells. The reason for such reduction and possible implications for the collective dynamics of cancer cells are discussed.

## INTRODUCTION

Understanding how cells go from a scattered distribution to a mechanically robust tissue on substrates and interfaces poses technological and theoretical challenges in the study of morphogenesis, tissue engineering, and cancer (1, 2). This emerging phenomenon is characterized by a transition from individual to collective motion where the interplay between the proliferation of cells and cell-cell interaction can lead to unusual non-equilibrium dynamics (3). Disruption on the regulation of one of these mechanisms can lead to pathologies with characteristic spatial patterns when compared to healthy tissues (4, 5). Therefore, it is key to control the morphology and mechanical robustness of the growing tissue, while guaranteeing a sustained delivery of nutrients to all cells (6, 7).

The capability of growing tissues on substrates is intrinsically connected with the ability of cells to migrate collectively and self-organize to generate spatial structures, with several mechanisms involved, such as cell-cell adhesion, cell motility and proliferation (8, 9, 10). Despite the impressive development of techniques for image capture and processing at the microscale, the identification of the relative importance of each mechanism is still a challenge. One promising strategy is to use techniques originally developed in the context of statistical mechanics and soft matter physics to characterize the structure and heterogeneous distribution of cells and tissues and their dependence on the experimental conditions (7, 11, 12, 13). For example, measurements of Moran’s I statistics, pair correlation function, and Ripley’s K-function are used to assess the degree of correlation on spatial distributions (14, 15, 16) and are instrumental to distinguish structural patterns of aggregation at various length scale, as in the case of benign and malign cancer cells (17). When combining experiments and in silico modeling, these measurements can help parametrizing the model parameters and thus infer the relative importance of each mechanism (3, 18, 19).

To better understand the underlying physical mechanisms of cell-tissue morphogenesis, a series of models have been developed. Examples include mean-field models (20, 21), the Fisher-Kolmogorov equation applied to the study cell migration (22), models based on cellular automata (23), vertex and Voronoi models (11, 24, 25, 26), cell dynamics based on Potts model or phase-field models (27, 28). Many models neglect the non-equilibrium dynamics and heterogeneity introduced by cell proliferation, an essential property, especially in the context of cancer (3, 29). Since the aim is to show how to infer the rate of each mechanism by combining in silico modeling with in vitro experiments, here, we develop a particle-based model to study the dynamics of adhesion, motility, and proliferation of cells on a flat substrate. *In vitro* experiments were performed using a culture of glioblastoma multiform (GBM) cell line, U87MG. The position of the cell nucleus was determined automatically with image processing algorithms and the time evolution of the spatial cell-cell correlation analyzed over a period of 24 h. By parametrizing the adhesion and proliferation rates in the model, it was possible to reproduce the evolution of the two-dimensional spatial heterogeneous distribution of cells, which provides insight into the underlying dynamics. The results revealed a reduction of cell-cell adhesion in response to the increase of cell density on the substrate as a function of time. This mechanism is consistent with a reduction in contact inhibition and consequently an enhancement on GBM cells’ migration, which in an *in vivo* scenario is translated usually in brain parenchyma invasion, hindering current local and regional therapies (30, 31).

## RESULTS

Glioblastoma multiforme (GBM) cells were cultured *in vitro* for 24 hours. The position of each cell on the flat substrate (bottom of the well) was identified by staining cells’ F-actin microfilaments of the cytoskeleton (red channel) and nuclei (blue channel) as depicted in Fig. 1(a). In the pictures obtained from this staining, the blue channel was separated to determine the position of each nucleus at each time, for 2, 4, 6, 8, 12, and 24 hours, as shown in Fig. 1(b). It was then traced a boundary in the blue channel to identify each nucleus (Fig. 1 (c)), which was used for subsequent analyses. To validate the image processing algorithm, in Fig. 1(d) is the evolution of the number of adhered cells as a function of time, which is in good agreement with the observed increase on dsDNA (see Fig. 1(e)).

**Fig. 1.**
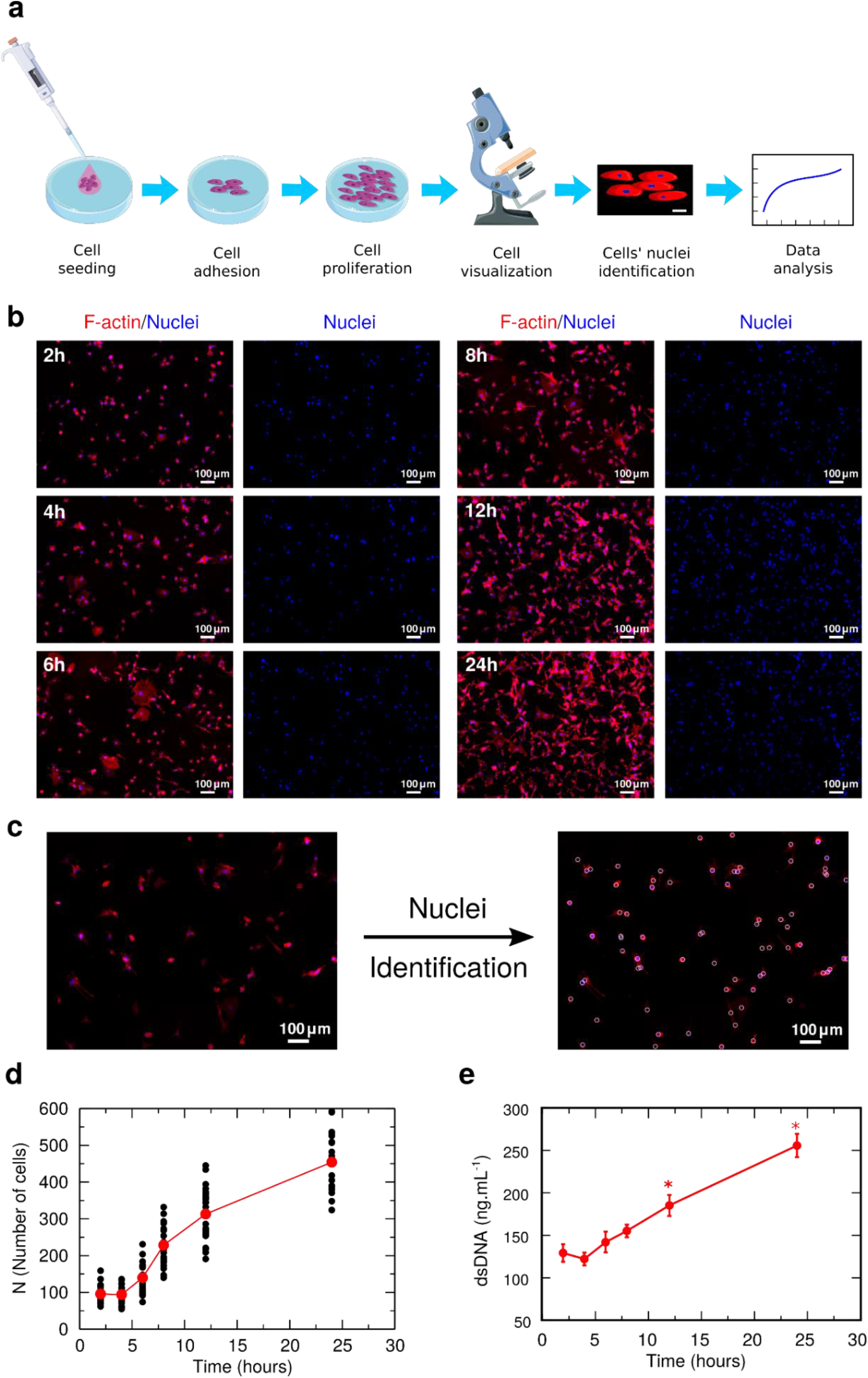
Cell culture and image processing. **(a)** Schematic representation of the culture of glioblastoma multiforme (GBM) cells on a substrate and posterior analysis using image processing. **(b)** Representative images of stained GBM cells for F-actin microfilaments of the cytoskeleton (red channel) and their nucleus (blue channel) at different instances of time. **(c)** Example of identification of each nucleus using image processing techniques, resolving the position of cells on the substrate after 6 hours of culture (Scale bar: 100 *μm*). From the identification of each nucleus, we quantify the proliferation as a function of time. **(d)** Evolution of the number of cells *N* as a function of time. Each black circle is the result for one image and the red circles are the average for each instant of time. To validate the image process algorithm, we show in **(e)** the dsDNA quantification indicating the number of cells adhered on the substrate along the 24 hours of culture. (*) denotes statistically significant differences (*p* < 0.05) comparing to 2 hours of culture. The similarity between these two measures indicates that the developed algorithm is correctly identifying the adhered cells.

To characterize the spatial distribution of nuclei in the substrate, it was used the Ripley’s K-function, which is defined as the average number of nuclei within a distance *r* of a randomly chosen nucleus rescaled by the average density. For a random distribution of nuclei (Poisson process) *K*(*r*) = *πr*^2^. Thus, when plotting *K*(*r*)/*r*^2^, any deviation from *π* indicates spatial correlations in the distribution of nuclei. Values above *π* would indicate a tendency to have more cells close to another cell (positive correlation), while values below *π* suggest cell depletion around a cell (negative correlation). Figure 2(a) shows K(r)/r^2^ as a function of the distance *r*, for different values of the average number of adhered cells ⟨*N*⟩ on the substrate. For a low number of GBM adhered cells, the average *K*(*r*)/*r*^2^ has a large deviation from a random case, but as the density increases, the average *K*(*r*)/*r*^2^ moves towards *π*. To understand the decrease in spatial correlation as a function of *N*, in Fig. 2(b) is the evolution of *K*(*r*)/*r*^2^ as a function of the number of cells *N* for a specific distance *r* = 20 *μm* (see Supporting Information Fig.S1 for the time evolution of *K*(*r*)/*r*^2^). As the number of cells adhered on the substrate increases, the correlation rapidly decreases converging to a value close to *π*, therefore indicating a decrease in spatial correlations.

**Fig. 2.**
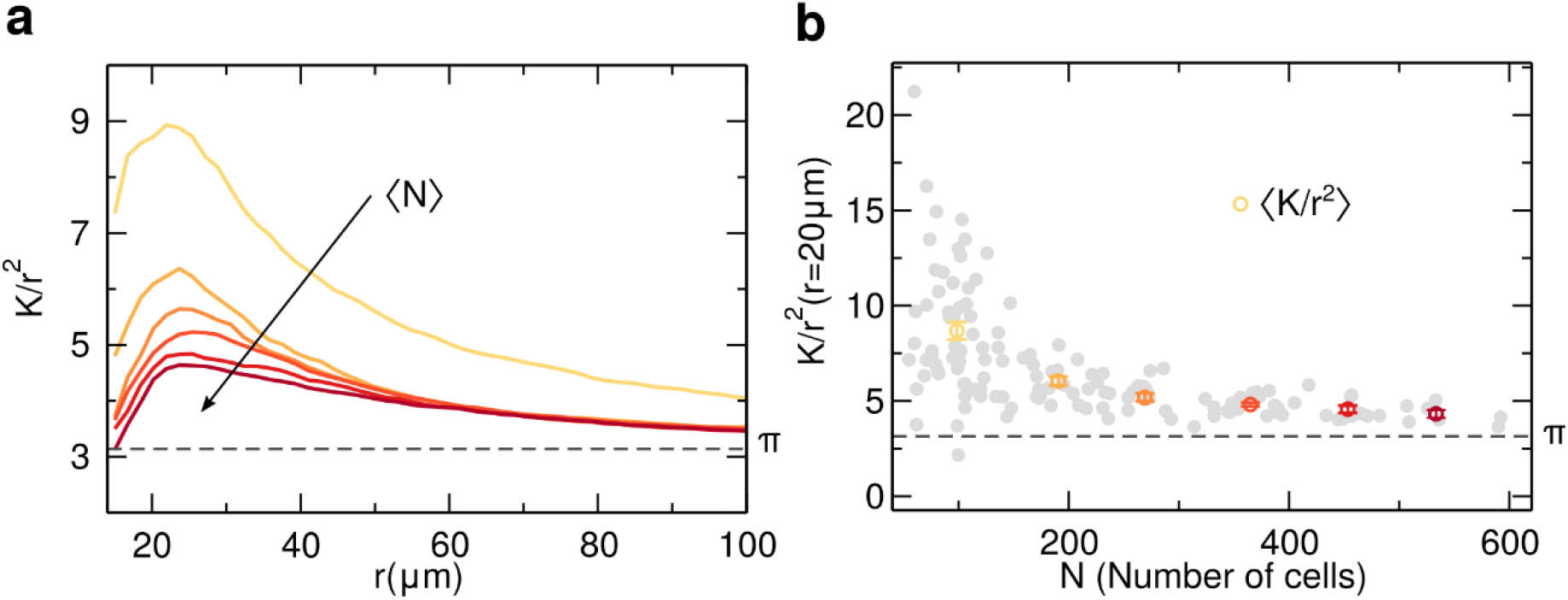
Proliferation and spatial correlations. Ripley’s K-function was used to characterize the distribution of the nuclei in the substrate. To enhance how far the experimental spatial distribution of glioblastoma multiform (GBM) cells is from a random process, we show *K*/*r*^2^ as a function of the distance *r* for values larger than 15 *μm*, avoiding imprecisions from a possible superposition of two nuclei. In **(a)** it is shown the average *K*/*r*^2^ for different values of the average number of cells in the substrate ⟨*N*⟩. The colors represent ⟨*N*⟩, from yellow (low density) to dark red (high density), following the linear spaced bins shown in (b). With a low density of cells in the substrate (⟨*N*⟩ = 98), it was observed that the distribution has a distinct peak and decays slowly to *π*, however, this correlation peak start to decrease as ⟨*N*⟩ increases during the 24 hours of experiment. In **(b)** it is depicted the evolution of the spatial correlation *K*/*r*^2^ as a function of the number of GBM cells (*N*) for a distance *r* = 20 *μm*. Each grey circle is a different measurement. The colored circles (yellow to dark red) are averages from equal spaced bins on *N*, exhibiting a rapidly monotonic decrease as the number of cells increases.

### Particle-based model

To model the proliferation of the GBM cells on a flat substrate, three mechanisms are considered: cell proliferation, motility, and cell-cell adhesion. Each GBM cell is represented by a circular particle with a radius of *r*_0_ = 10 *μm*, 40% larger than the average radius of a GBM cell nucleus (see Supporting Information Fig. S2 for the distribution of nucleus radius). This value was considered to simulate what would be a minimum distance of interaction between cells due to an effective shape and, as shown in the Supporting Information Figure S5, the results are robust to small variations in this value for the particle radius. In the experiments, each well is prepared with an initial quantity of 10^4^ cells and have a total substrate area of 1.93 *cm*^2^. Since each experimental image has an area of about 10^−2^ *cm*^2^ and assuming that the deposition of cells occurs before two hours, we estimate an average initial number of cells *per* image of about 50 cells. Therefore, in silico, each sample starts with 50 particles distributed uniformly at random on the substrate.

Cells divide at a rate λ: an offspring cell is positioned at random, at a distance *d* = 2*r*_0_ from the original one, and the division is only successful if the offspring cell does not overlap any other cell. To model cell motility, a Brownian dynamics was implemented with a diffusion coefficient τ*D*, where *D* is the diffusion coefficient of a cell and τ is an adimensional constant that mimics the effect of the cell-cell adhesion, which usually results in a decrease in cell motility. If a cell has another one at a distance lower than 2*r*_0_, τ ≠ 1, otherwise, τ is unitary (see model scheme in Fig. 3(a)). A similar model of interaction has been used successfully to describe the formations of patterns in extra-cellular matrix by the collective migration of fibroblasts in Refs. (8) and (32). The discrete equations of motion for the coordinates *xi* and *yi* of each cell *i* are:

**Fig. 3.**
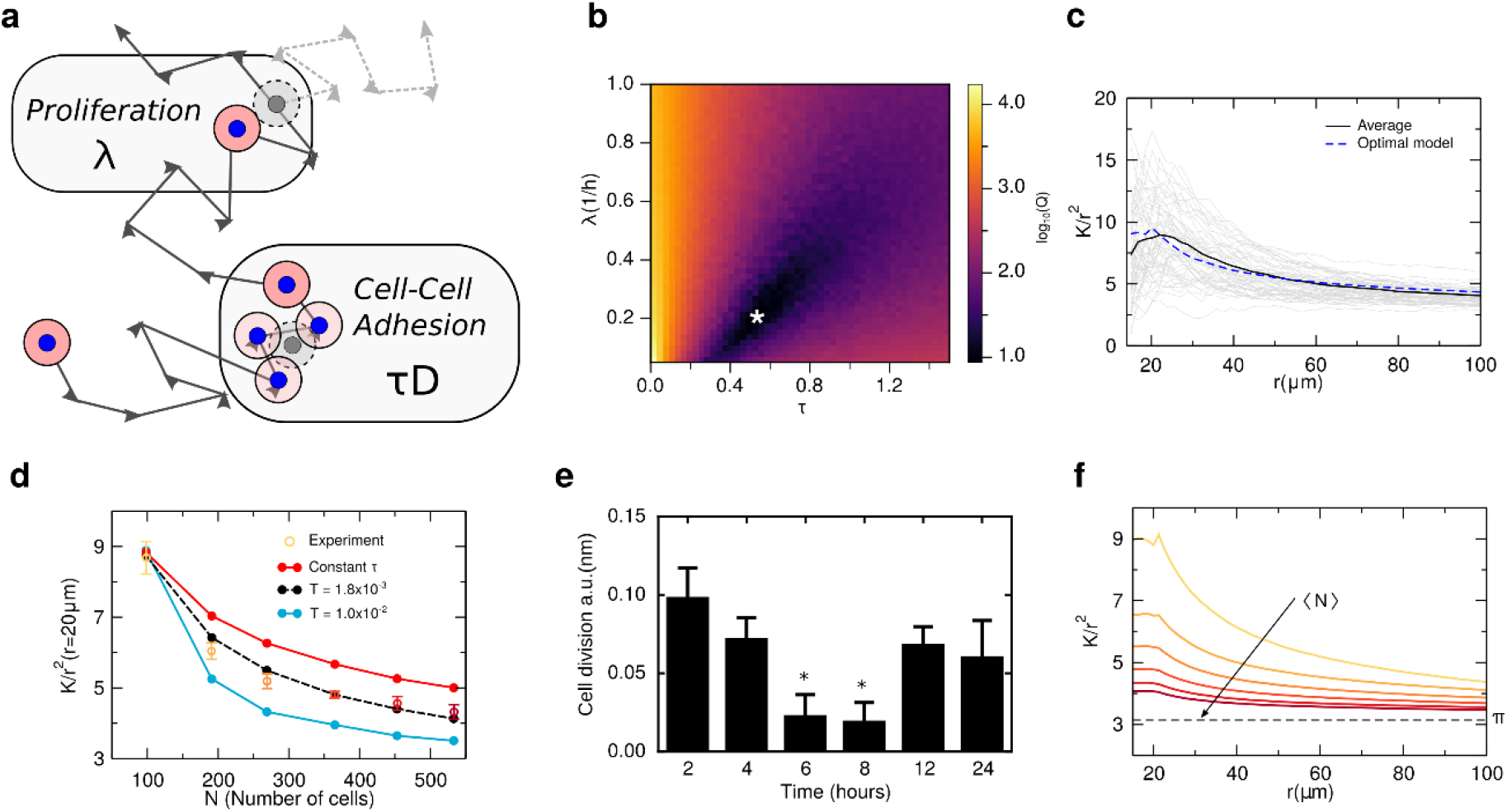
Calculation of the proliferation rate and cell-cell adhesion. **(a)** Schematic representation of the dynamics of a cell modeled in silico. Each cell performs a Brownian motion with a diffusion coefficient *D*. To account for cell-cell adhesion, when two cells overlap the diffusion coefficient is changed to τ*D*, with τ ≠ 1.0. As show in the scheme, the proliferation is controlled by the parameter λ, that sets the rate of cell division (gray circle). In **(b)** is the logarithm of the quadratic difference *Q* between the experiment and simulation averages over 200 independent runs for each pair of λ and τ. We observe that Q is minimized for *λ*_0_ = 0.2 and τ_0_ = 0.52 (white star). In **(c)** is a comparison between the experimental average *K*(*r*)/*r*^2^ (black line) and the model using the optimal parameters (dashed blue line). Each gray line refers to a sample from the experiment within that range of *N*. **(d)** Average of *K*/*r*^2^(*r* = 20 *μm*), as a function of *N*. We assume that τ is a function of the density, with the form given by Eq. (3). For *T* = 0.0 (red connected circles) the simulations systematically overestimate the experimental values (yellow to dark red circles). However, using *T* = 1.8 × 10^−3^ we obtained a quantitative agreement between simulations and *in vitro* experiments (black connected circles). Each point from the simulation is an average over 10^3^ samples. **(e)** The proliferation rate of GBM cells was assessed experimentally using a colorimetric immunoassay based on the measurement of BrdU incorporation during DNA synthesis along the 24 hours of culture. (*) denotes statistically significant differences (p < 0.05) comparing to 2 hours of culture. **(f)** Average *K*(*r*)/*r*^2^ as a function of *r* for different values of *N*, the same as the average number of GBM cells in the *in vitro* experiment, and using *T* = 1.8 × 10^−3^. Each curve is a result of 10^3^ simulations and the colors represent the average number of cells following the linear bins shown in Fig. 2(b).

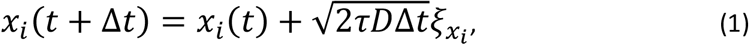

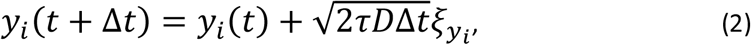

where 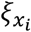 and 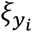 are noise term, following a Gaussian distribution with average zero and unitary dispersion and Δ*t* is the time step. See Supporting Information Figure S3 for more details about the numerical implementation of the model.

### Inferring the values of the rates

We fixed the diffusion coefficient to *D* = 500*μm*^2^/*h*, estimated by previous experimental measures for GBM cells (33). To adjust λ and τ, several simulations were performed for a range of values of *λ ∈* [0.15, 1.0], in units of inverse of hour, and τ *∈* [0.0, 1.5], until the number of particles reaches *N* = *N*_0_, the same average number of cells for the first measurement in the experiment (*N*_0_ = 98). For each pair of parameters, using 200 independent simulations, the average of *K* (*r*)/*r*^2^ was calculated, as well as the quadratic deviation 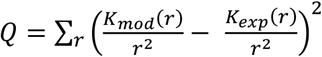 to the experimental values. Figure 3(b) shows that there is a set of λ and τ That minimizes the deviation to the experiment with a low density of cells. Using the optimal values of the parameters, λ_0_ = 0.2 and *τ*_0_ = 0.52, the average of K_mod_(r)/r^2^ is the one shown in Fig.

3(c) (blue dashed line). It is in good quantitative agreement with the values obtained for the *in vitro* experiments (black line).

### Density dependent cell-cell adhesion

The Ripley’s K-function at *r* = 20 *μm* for the values of *λ*_0_ and τ_0_ inferred from the results for *N* = 98 are represented by the red circles in Fig. 3(d). It is clear that with constant values of *λ*_0_ and τ_0_, the particle-based model systematically overestimates the spatial correlations as a function of *N*. Measures of the BrdU incorporation during DNA synthesis along the 24 h of culture, shown in Fig. 3(e), suggest a constant rate of proliferation, thus λ should be constant. Thus, it is hypothesized that τ depends on *N*. To consider such dependence, it was assumed an exponential decay,

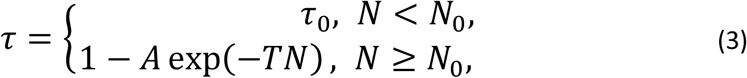

Where *A* = (1 − τ0) exp(*N*0*T*) is a constant and the threshold of *N*0 is the average number of cells considered to infer τ_0_. From Eq. (3) τ converges to unity for large values of *N*. The dependence on *N* of τ is parametrized by a new parameter *T*, which sets a characteristic density. For *T* = 0.0, the results for constant τ are recovered.

Results for different values of *T* are shown in Figs. 3(d). From the simulations, we estimate *T* = 1.8 × 10^−2^ for the *in vitro* experiments (see Supporting Information Fig. S4 for a detailed calculation of the optimal *T*), and the correlations for all values of *r* are shown in Fig. 3(f) in good agreement with the *in vitro* experiment (see Fig. 2(a)). This finding suggests that the GBM cells are adapting their interactions as they proliferate, for example, through the formation of multilayer structures.

## DISCUSSION

GBM is very common cancer that presents an extraordinarily bad prognosis, resulting into a very low survival rate (≈ 15 months) (34). One of GBM hallmarks is its propensity to metastasize to different sites within the brain. Unfortunately, effective therapies capable to target migrating cancer cells that origin metastasis are still not available (35). It has been shown that the interaction between cancer cells is a crucial step to promote cell proliferation, survival and ultimately metastasize (36). In fact, it is described that during metastization, cancer cells suffer a transformation known as epithelial to mesenchymal transition that results in a decrease of cell adhesion and polarity, potentiating cell migration and proliferation (37). But, a lot is still to be understood on the collective dynamics of GBM cells.

To gain insight into the physiology and dynamics of GBM cells proliferation on a flat substrate, we developed a minimum model that incorporates three key mechanisms: cell motility, cell-cell adhesion, and proliferation. We show that is possible to parametrize the rates associate with each one of these mechanisms to reproduce the collective dynamics observed in an *in vitro* experiment. The experiment was performed using a culture of glioblastoma multiform (GBM) cell line. From 24 hours experiment, a series of pictures of stained cells were obtained and, using image processing, the position of cells was determined. The time evolution of the two-dimensional spatial cell-cell correlation as a function of cell density in the substrate was analyzed. We demonstrated that by controlling the GBM cell-cell adhesion and proliferation rates in the model it is possible to explain the observed cell-cell spatial correlations providing insights into the underlying mechanisms. The simulations indicate that GBM cell-cell adhesion must decrease when increasing cell density, allowing the cells to move over each other more easily. This decrease in cell-cell adhesion might be related to the absence of contact inhibition in cancer cells (30, 38). In the special case of GBM, it is described that cells proliferate while there are enough nutrients and oxygen (39). But, when there is a deficiency of one, cells are prompted to migrate. Since proliferation reach a plateau (Fig. 1), it may indicate that GBM cells are migrating over each other looking for a more favorable environment within the flat substrate.

In conclusion, there is a growing interest on combining in silico modeling with experiments to provide insight into underlying mechanics of cellular systems (40,41,42). Exemplarily, we have shown that even with a simple, particle-based model is possible to infer the rate of key mechanisms and find new behavior. In particular, a parametrization of the model to recover the experimental results revealed a dependence of cell-cell adhesion with cell density, result consistent with experimental observations of epithelial to mesenchymal transition (43).

## MATERIALS AND METHODS

### *In vitro* cell culture

Glioblastoma multiform (GBM) cell line, U87MG was cultured in DMEM, (Sigma) supplemented with 10% fetal bovine serum (Alfagene) and 1% penicillin and streptomycin (Thermo Fisher Scientific) at 37°C under a controlled humidified atmosphere containing 5% CO2. The medium was changed twice a week. When the confluence of cells reached ≈ 80%, they were passaged and plated at an initial density of 10^4^ cells per well in a 24-well plate. At different times (0, 2, 4, 6, 8, 12, and 24 hours), the proliferation of cells and morphology were assessed.

### dsDNA quantification of the GBM cells

The total double-stranded DNA (dsDNA) quantification was analyzed at each time to assess the number of cells adhered to the substrate. For that, ultrapure water was added to each well, and cells were incubated for 1 hour at 37°C and stored at −80°C until analyzes. GBM cells in ultrapure water were then sonicated for 10 minutes and used for dsDNA quantification using the Quant-iT PicoGreen dsDNA kit (Molecular Probes, Invitrogen), according to manufacturer’s instructions. Briefly, samples were transferred to a 96-well white plate and diluted in TE buffer. After adding the Quant-iT PicoGreen dsDNA reagent, samples were incubated for 10 minutes at RT in the dark, and fluorescence was quantified using a microplate reader (Biotek Synergy HT) with Ex/Em at 480/530 nm. RFUs were converted into ng.mL^-1^ using a standard curve of DNA in the range of 1 - 2000 ng.mL^-1^.

### Position identification of the GBM cells

At each time aforementioned, the position of the GBM cells was identified by staining cells for F-actin microfilaments of the cytoskeleton and their nuclei. For that, cells were washed with phosphate buffer saline (PBS, Sigma-Aldrich), fixed with 10% Neutral Buffered Formalin (ThermoFisher Scientific) for 20 minutes and permeabilized for 5 minutes with 0.1% v/v Triton X-100 (Sigma-Aldrich) in PBS. Afterward, F-actin filaments were stained with Phalloidin– Tetramethylrhodamine B isothiocyanate (Sigma-Aldrich, 1:100), and the nuclei were counterstained with 1:1000 of the stock of 4,6-Diamidino-2-phenyindole, dilactate solution (DAPI, 1 mg/mL, Biotium). Finally, stained cells were observed under fluorescence microscopy (Zeiss).

### Proliferation of the GBM cells

To assess the proliferation of the GBM cells, a colorimetric immunoassay based on the measurement of BrdU incorporation during DNA synthesis named Cell Proliferation ELISA, BrdU (Roche, Laborspirit) was used according to manufacturer’s instructions. Briefly, 2 hours before each time, GBM cells were labeled with 100 μM of BrdU. Then, at each time, cells were rinsed with PBS, dried at 60°C for 1 hour, and stored at 4°C until the end of the experiment. Then, the reagent FixDenat was added to the cells and incubated for 30 minutes at room temperature. After removing FixDenat, anti-BrdU-POD was added and incubated for 90 minutes at room temperature. Finally, cells were washed three times with PBS, incubated with substrate solution for 30 minutes at room temperature, the reaction was stopped with 1 M H2SO4, and absorbance was measured at 450 nm using a microplate reader (Biotek Synergy HT).

### Statistical analysis

For *in vitro* cell culture experiments, statistical analyses were performed using GraphPad Prism 6.0 software. For the analysis of GBM cells’ dsDNA quantification and proliferation rate, the non-parametric Mann–Whitney test was used to compare two groups, whereas the comparison between more than two groups was performed using the Kruskal–Wallis test, followed by Dunn’s comparison test. Concerning cumulative proliferation of cells, the non-parametric Friedman test was used followed by Dunn’s comparison test. The critical level of statistical significance chosen was *p* < 0.05.

## Acknowledgments

The authors acknowledge financial support from the Portuguese Foundation for Science and Technology (FCT) under Contracts no. PTDC/FIS-MAC/28146/2017 (LISBOA-01-0145-FEDER-028146), UIDB/00618/2020, and UIDP/00618/2020. F.R.M. also acknowledges FCT for her contract under the Transitional Rule DL 57/2016 (CTTI-57/18-I3BS(5)).

