## Supplemental Text for "Combining experiments and in silico modeling to infer the role of adhesion and proliferation on the collective dynamics of cells"

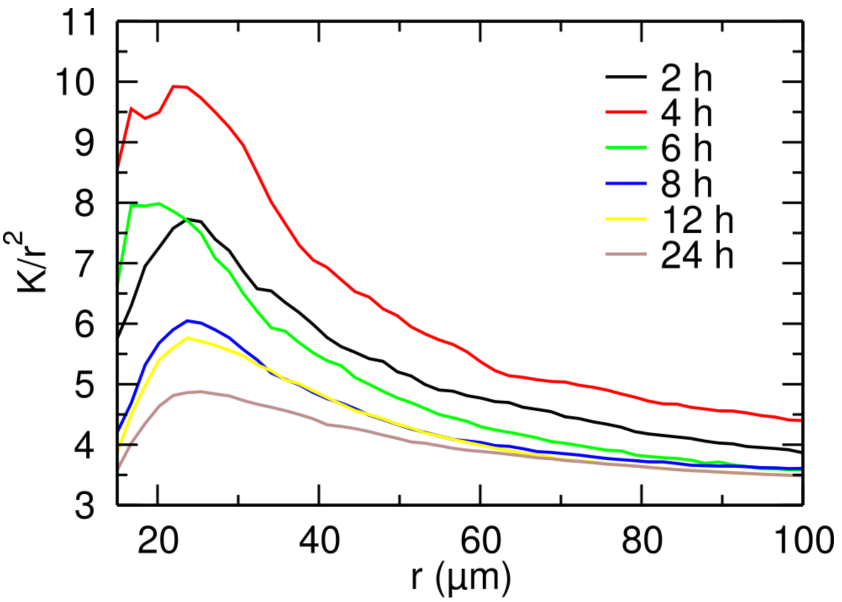


**Fig. S1. Spatial correlations stratified by time.** Average of $K/r^2$ as a function of $r$ for each time instant using the *in vitro* experimental data. Since the number of GBM cells in the substrate increases with time, we expected to see a similar behavior as in Fig. 3(A). We see some deviations for the first instants.

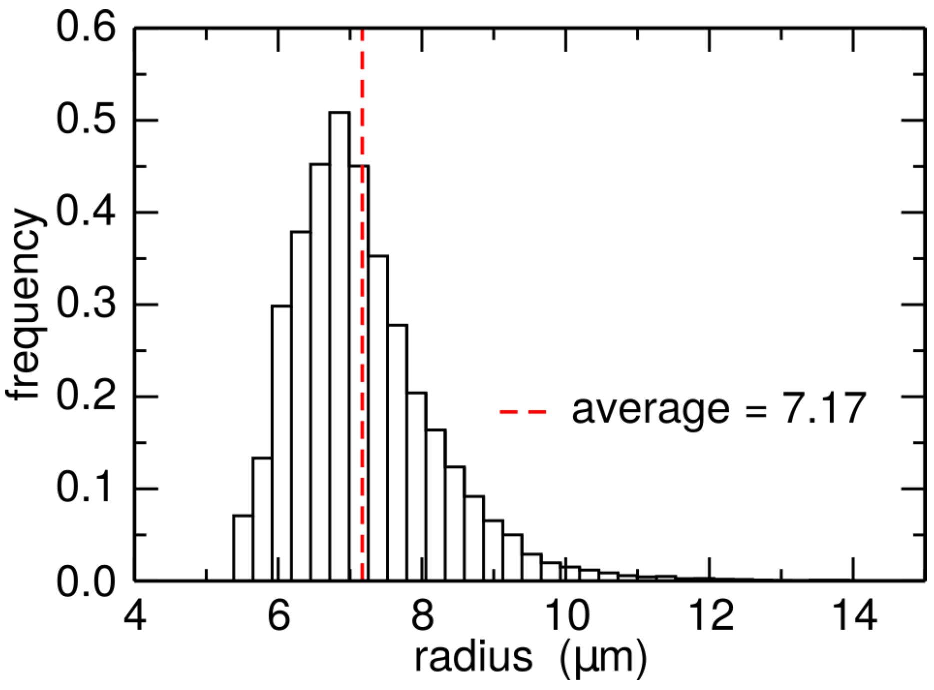


**Fig. S2. Nucleus size distribution.** Normalized histogram for the radius of each cell, measured with the image processing algorithm, assuming a circular shape. Results were obtained considering all pictures along the 24 h experiment. These results suggest a narrow distribution and an average of 7.17 $\mu m$.

### Model Dynamics

In Fig. S3 are the dynamical properties for different sets of parameters. In Fig. S3(A) is the time dependence of the number of cells $N$. If $\tau \neq 0$ the cells can move on top of each other, creating new gaps to fit new cells by division, allowing an unbounded growth of $N$. On inset of Fig. S3(A), we see that $\lambda$ controls how fast $N$ saturates, however, after the saturation, the growth rate will be controlled by $\tau$ and independent of $\lambda$.

From the simulations, it was possible to calculate the spatial correlations using $K(r)/r^2$ when reaching a specific value of $N$. We show in Fig. S3(B) that for $\lambda = 10$ and $\tau = 10^{-3}$ when the system reaches $N = 100$, the $K(r)/r^2$ depicts a characteristic peak around $r = 2r_0$, similar to the behavior found on the experiments (Fig. 3(A)). When increasing $N$ we also verify that $K(r)/r^2$ starts to decrease, a similar behavior to the *in vitro* experiments.

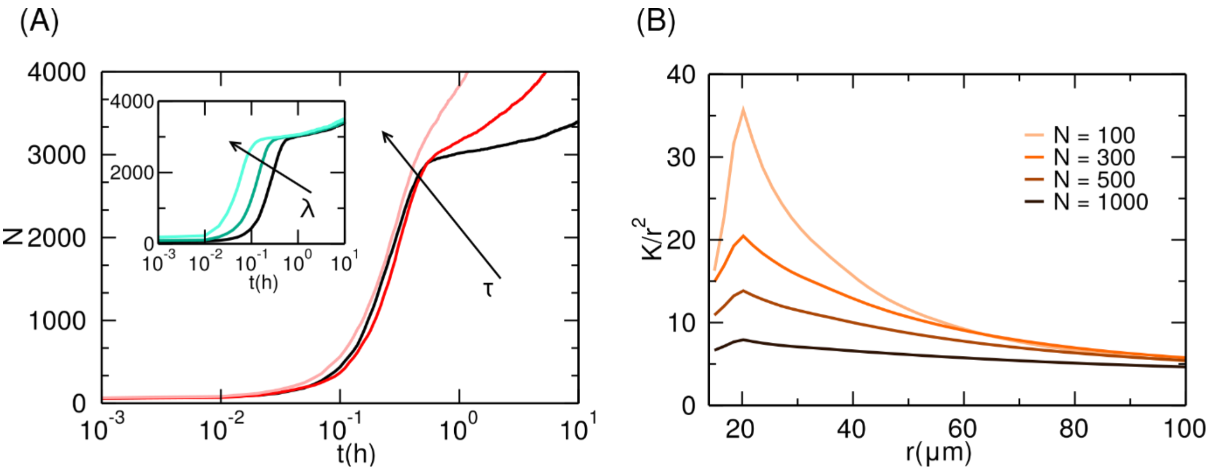


**Fig. S3. Model dynamics and spatial correlations.** Starting with 50 randomly positioned particles, we studied the time evolution of the number of particles $N\ (t)$ for different values of the proliferation rate λ and cell-cell adhesion parameter $\tau$. In (A) are the results for $\lambda = 10^2$, in units of inverse hour. Initially, $N$ grows rapidly, but saturates for a specific N, independent of $\tau$, the growth rate slows down. Three values for cell-cell adhesion were considered, namely, $\tau = 10^{-3}$ , $10^{-2}$, and $10^{-1}$. The growth rate in the slower regime depends on $\tau$. In the model, there is no upper limit for the number of particles since, for $\tau \neq 0$, it is possible to have cells overlapping and thus, there is always the possibility of forming gaps where cell division can occur. On the inset, $\tau = 10^{-3}$ and different values of $\lambda$ are considered, namely, $\lambda = 10^2$, $2 \times 10^2$, and $5 \times 10^2$, all in units of inverse hour. The duplication rate controls the growth rate in the first regime. In (B) is the spatial correlations measured by $K(r)/r^2$ for $\lambda = 10$ and $\tau\ =\ 10^{-2}$.

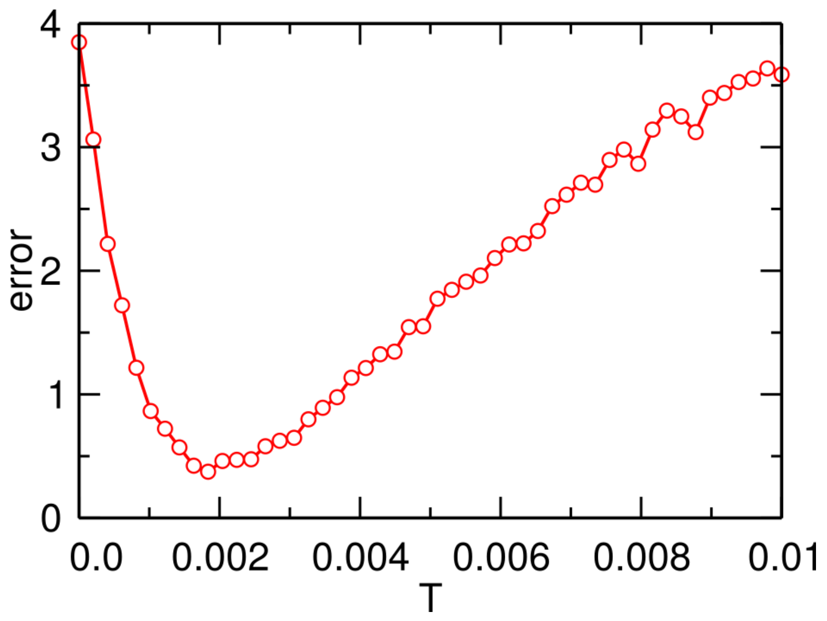


**Fig. S4. Determination of the parameter T.** From the simulations using Eq. (3) to update the GBM cell-cell adhesion $\tau$, we considered different values of $T$, and compared the numerical results for the spatial correlation for $r = 20\ \mu m$ to the one measured experimentally. We define as the *error*, the quadratic difference between the correlations from simulations and experiments. We see that there is a value of $T$ that minimizes this *error*, and for $T = 1.8 \times 10^{-3}$ our model shows the best agreement with the *in vitro* experiment indicating that $\tau$ have to decrease with the number of GBM cells.

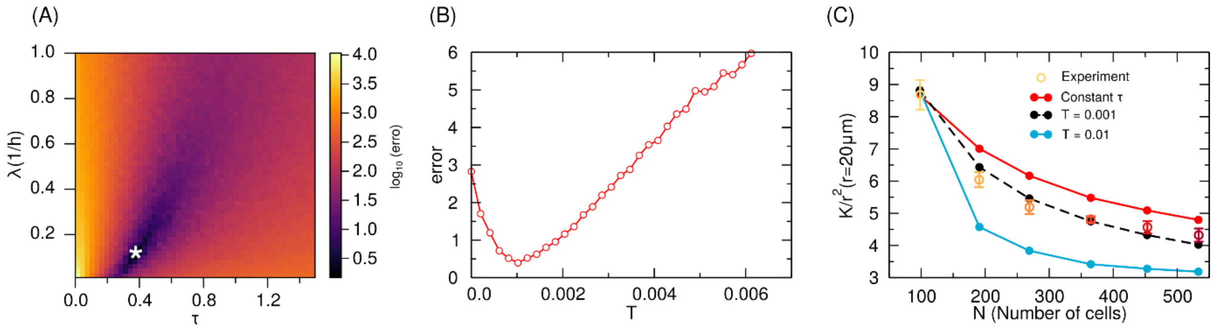


**Fig. S5. Results with different $r_0$.** The particle radius $r_0$ sets an effective minimum radius of interaction between two cells. Here, we show that even using a larger radius of $r_0 = 13\ \mu m$, the results are qualitatively the same. In (A), it is the colormap for the logarithm of the difference between the model and experimental spatial correlations using 200 simulations for each pair of $\lambda$ and $\tau$. It was observed that there is an optimal combination of $\lambda$ and $\tau$ where the difference is minimized (white star). The optimal values are $\lambda_0 = 0.1$ and $\tau_0 = 0.36$. Using these values and Eq. (2) for the evolution of $\tau$, it is shown in (B) that there is a value $T$ where the error is minimized. In (C) is the dependence on the number of cells N of $K/r^2(r = 20\ \mu m)$ for different values of $T$. Therefore, the indication of reduction of GBM cell-cell adhesion is robust to a change of $r_0$ in the model.
